# WormID-Bench: A Benchmark for Whole-Brain Activity Extraction in *C. elegans*

**DOI:** 10.1101/2025.01.06.631621

**Authors:** Jason Adhinarta, Jizheng Dong, Tianxiao He, Junxiang Huang, Daniel Sprague, Jia Wan, Hyun Jee Lee, Zikai Yu, Peng Liu, Hang Lu, Eviatar Yemini, Saul Kato, Erdem Varol, Donglai Wei

## Abstract

The nematode *C. elegans* is a premier model organism for studying neural circuit function due to its fully mapped connectome and genetically identifiable neurons. Recent advances in 3D light microscopy and fluorescent protein tagging have enabled whole-brain imaging at single-neuron resolution. However, extracting meaningful neural dynamics from these high-resolution recordings requires addressing three fundamental challenges: (i) accurate detection of individual neurons in fluorescence images, (ii) precise identification of neuron classes based on anatomical and colorimetric cues, and (iii) robust tracking of neurons over time in calcium imaging videos. To systematically evaluate these challenges, we introduce WormID-Bench, a large-scale, multi-laboratory dataset comprising 118 worms from five distinct research groups, along with standardized evaluation metrics for detection, identification, and tracking. Our benchmark reveals that existing computational approaches show substantial room for improvement in sensitivity, specificity, and generalization across diverse experimental conditions. By providing an open and reproducible benchmarking framework^1^, WormID-Bench aims to accelerate the development of high-throughput and scalable computational tools for whole-brain neural dynamics extraction in *C. elegans*, setting the stage for broader advancements in functional connectomics.

## 1 Introduction

The ability to robustly resolve whole-brain activity at single-neuron resolution remains a major challenge in neuroscience. *Caenorhabditis elegans (C. elegans*) serves as a powerful model system due to its fully mapped connectome and well-characterized neuronal classes [24,25]. Recent advances in 3D light-sheet microscopy and fluorescent genetic labeling (NeuroPAL) [27] enable the tracking of neuronal activity across the entire nervous system. However, a key bottleneck is the development of reliable computational methods to identify the activity for each neuron, which involves neuron detection and identification from multichannel NeuroPAL images, and neuron tracking from calcium images across diverse experimental conditions.

Automating these tasks has seen limited success; the dense distribution of similar-colored neurons and non cellular structure makes detection and identification challenging. And neural dynamics can make accurate tracking difficult, as the brightness and appearance of neurons can vary widely over time based on their firing activity [13]. Worse still, the imaging data is diverse in image format and appearance, biological variability, and batch effects across different labs and experimental setups [19,20,21,12,11,26]. Recently WormID [17] curated several publicly available datasets with complete or semi-complete ground truth annotations that span multiple imaging setups and laboratories. However, there remains a lack of clearly defined tasks, standardized evaluation metrics, and benchmarks for state-of-the-art computer vision methods in worm neural dynamics extraction.

To address this, we introduce WormID-Benc to evaluate whole-brain neural dynamics extraction in *C. elegans* in Fig. 1 . Our benchmark provides (1) a structured evaluation protocol for neuron detection, identification, and tracking; (2) reproducible metrics and ranking procedures to ensure fair and unbiased model evaluation; (3) comprehensive benchmark results on the WormID dataset.

**Fig. 1.**
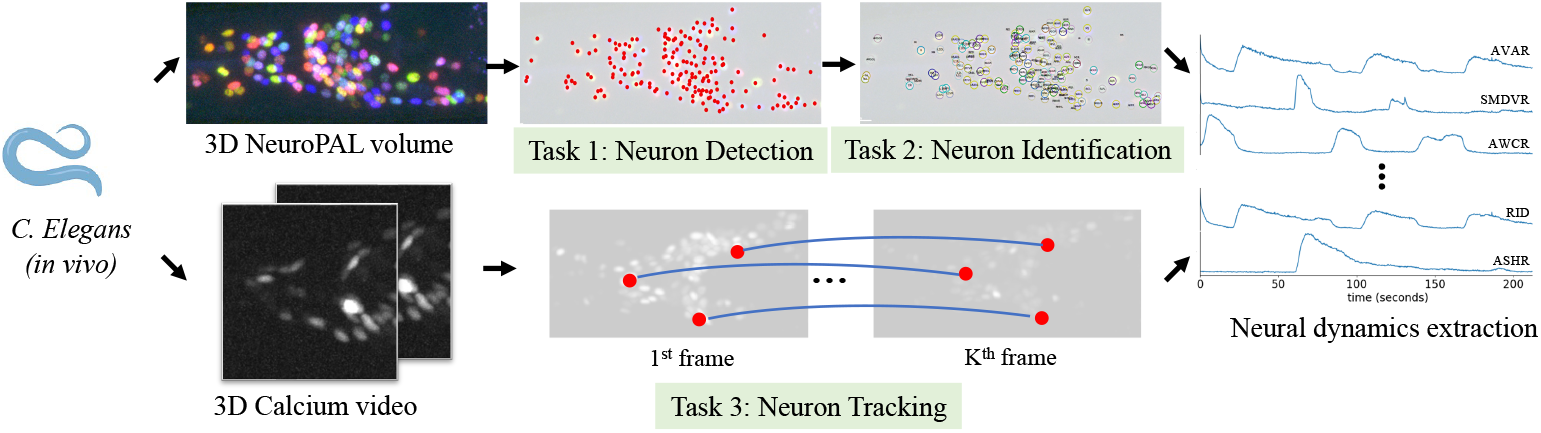
WormID-Bench. We benchmark each step of the computational pipeline to extract neural dynamics of *in vivo C. elegans* from microscopy images.

## 2 Related Work

We limit our review to light microscopy which is standard for *in vivo* live imaging.

### 3D Cell Detection Benchmarks

3D cell detection is always the first step for any neuron-level imaging-related task. Previously Alwes *et al*. [1] collected and annotated volumes of 3D images of histone fluorescent protein expression in Parhyale. Another valuable resource is jGCaMP8 transgenic mice dataset, which applies two-photon imaging to record neurons in mouse visual cortex [22]. However, there are few large-scale datasets that cover neuronal volumes in *C. elegans* [14] with comprehensive annotations for cell detection, identification, and cell tracking. Unlike previous datasets, *C. elegans* microscopy images tend to have lower resolution and have a more non-cellular structure, making the segmentation task challenging.

### 3D Neuron Identification Benchmarks

Neuron identification from fluorescent microscopy images is notoriously difficult. While NeuroPAL method [27] deterministically colors every neuron a stereotyped fluorescent barcode in *C. elegans*, which is identical across all stages of development for all individuals. The color code and relatively fixed position between neurons greatly simplify the identification problem by reducing the number of potential labels for a neuron.

### 3D Cell Tracking Benchmarks

The popular Cell Tracking Challenge [8] consists of ten 3D time-lapse microscopy datasets with various non-neuron cells. It provides dense segmentation annotations at each time frame, along with tracking annotations that indicate correspondences between frames and cell-splitting events. For neural dynamics extraction, annotating cell center points is sufficient, while minimizing significant manual labeling labor. To our knowledge, the 36 worms video dataset we are using is the only dataset with annotations for neuron subtype identities, their tracked positions, and their neural activity.

## 3 Dataset

For this benchmark, we use the WormID corpus of NeuroPAL and calcium imaging datasets curated in Sprague *et al*. [17], consisting of seven datasets from five different labs comprised of 118 total worms (Tab. 1). Further details can be found at WormID.org and in Sprague *et al*. [17].

**Table 1.** Summary of WormID-Bench datasets and their parameters. #DET, #ID, and #TRAJ represent the average number of annotated neuron for detection, identification, and trajectory across samples respectively.

| Dataset | resolution<br>($\mu\text{m}/\text{px}$) | 3D NeuroPAL | | | 3D Calcium Video | | |
| --- | --- | --- | --- | --- | --- | --- | --- |
|  |  | #samples | #DET | #ID | #samples | #frames | #TRAJ |
| NP [27] | $0.21 \times 0.21 \times 0.75$ | 10 | 193 | 190 | × | × | × |
| HL [5] | $0.33 \times 0.33 \times 1.0$ | 9 | 119 | 64 | × | × | × |
| SF [3] | $0.54 \times 0.54 \times 0.54$ | 38 | 70 | 70 | × | × | × |
| EY [12] | $0.27 \times 0.27 \times 1.5$ | 21 | 177 | 175 | 21 | 961 | 177 |
| KK [23] | $0.32 \times 0.32 \times 1.5$ | 9 | 154 | 154 | 9 | 1646 | 155 |
| SK1 [17] | $0.16 \times 0.16 \times 1^1$ | 21 | 111 | 48 | 4 | 1500 | 108 |
| SK2 [17] | $0.32 \times 0.32 \times 0.75^1$ | 10 | 173 | 49 | 2 | 3000 | 131 |

### Data Selection

For the detection and identification (ID) tasks, we omit 14 worms that contain significant non-linear deformities or color artifacts since many of the models we test assume that inputs are roughly aligned along the principal axes of the worm body and rely on color information. This leaves 104 remaining datasets that we use to train and benchmark each of the detection and ID approaches. For the tracking task, we omit 38 worm videos in SF [3] which does not provide enough information to extract ground truth annotations.

### Data Split

Most previous approaches to the tasks outlined here typically only train and test on a small subset of data, usually all collected by one lab, limiting the generalizability of these approaches. We split the datasets into five equal groups with balanced representation from each dataset. For each task, we perform 5-fold cross-validation and report results across multiple metrics. For the ID tasks, the whole NP dataset, which is the standard reference dataset for NeuroPAL imaging, is used for training all groups but is not used in evaluation. Thus, we report an average of cross-validated metrics across 104 worms for the detection task, 94 worms for the ID task, and 36 worms for the tracking task.

## 4 Benchmark

### 4.1 Task 1: 3D Neuron Detection in 3D NeuroPAL Volumes

#### Objective

The first step is to detect neurons in the NeuroPAL volumes while distinguishing them from non-neural cells and structures of the nematode. Due to the lack of visible cell boundaries and the low-resolution of the images, each neuron was annotated with the center point instead of the mask. This is particularly challenging due to the presence of background artifacts, variability in fluorescence intensity, and differences in imaging setups across laboratories.

#### Evaluation Metrics

We use evaluation metrics from the OCELOT Cell Detection Challenge [16], primarily the mean F1-score, along with precision and recall. Since our neuron annotations are point-based, we introduce a distance threshold to assess detection accuracy. A detection is considered correct if it falls within distance *d*_th_ of a ground truth neuron; otherwise it is a false positive. If multiple ground truth neurons are within *d*_th_, the nearest one is matched. To account for varying proximity requirements, we evaluate at two distance thresholds: *d*_th_ ∈{3*μm*, 6*μm*} , roughly 1-2 times the nucleus diameter, which ensures spatial accuracy and alignment with calcium imaging for cell tracking.

#### Baseline Models

Neuron detection methods can be categorized into point-based and mask-based approaches. Point-based methods predict a heatmap of neuron center locations, with peak detection algorithms used to extract precise coordinates, such as nn-UNet adopted for heatmap regression [6]. Mask-based methods leverage pre-trained generalist models, such as CellPose [18] and microSAM [2], to segment neuron regions. The centroid of each masked object is extracted as its cell coordinate.

### 4.2 Task 2: 3D Neuron Identification in 3D NeuroPAL Volumes

#### Objective

The 302-neuron nervous system of *C. elegans* is eutelic, meaning each neuron has a consistent identity across individuals. The computational task involves the 302-way classification for each detected neuron center in a 3D NeuroPAL image volume. Despite its stereotyped nervous system, neuron identification is challenging due to natural variability in cell positions for different individuals, body distortion during movement, color overlap for several neurons, and imaging inconsistencies between different labs. Moreover, the input detection results can have missed neurons or false positives, and some neurons remain difficult to distinguish even for experts.

#### Evaluation Metrics

We adopted the standard classification accuracy to measure the proportion of correctly labeled neuron centers in test images. The accuracy is reported for both top-1 and top-5 ranked assignments. Based on the number of neurons labeled, datasets are categorized into low and high label counts, as fewer labels typically yield higher accuracy since the easiest neurons are identified first. In contrast, fully labeled datasets tend to have lower accuracy due to the ambiguity of harder-to-identify neurons.

#### Baseline Models

Neuron identification methods fall into two main categories: alignment-based and classification-based approaches. Alignment-based methods perform non-rigid point cloud registration to align predicted neuron positions with a reference template. Techniques such as Coherent Point Drift (CPD) [10] are commonly used, followed by neuron label assignment using the Hungarian algorithm [7] or a learned statistical atlas [21]. Classification-based methods train machine learning models to directly predict neuron identities. Examples include a transformer-based approach [28] and a graph-based conditional random fields model [5]. For learning-based methods, we evaluate both pre-trained and fine-tuned models to assess their generalization across datasets.

### 4.3 Task 3: 3D Neuron Tracking in 3D Calcium Videos

#### Objective

The task involves tracking neurons across video frames and estimating activity from image intensity. Neuron tracking is challenging due to biological noise, deformations, missing neurons, and variability in data collection across labs. Previous methods have used pose registration to align neurons under deformation of the worm body. Additionally, inactive neurons may be undetectable in the calcium images while remaining visible under fluorescent channels invariant to excitation.

#### Evaluation metrics

We adopted the evaluation metrics from the Cell Tracking Challenge [8], specifically Detection Accuracy (DET) and Tracking Accuracy (TRA). TRA is based on the Acyclic Oriented Graph Matching measure [9], which computes the edit-distance between graphs, robustly handling cases where tracks are split or swapped. As in Task 1, setting a distance threshold is crucial to assessing detection and tracking accuracy. Therefore, we apply two distance thresholds, *d*_th_ ∈{3*μm*, 6*μm*} .

#### Baseline methods

Neuron tracking methods can be classified into matching-based and propagation-based approaches. Matching-based methods detect neuron center points in individual video frames and establish correspondences across frames. Matches are determined using integer linear programming (ILP) [4] or directly predicted by a feed-forward neural network. Propagation-based methods estimate optical flow to compute dense voxel-level correspondences between neighboring frames. Neuron center points are then propagated from the previous frame to the current frame by following these correspondences [15].

## 5 Results

For each task, we report 5-fold cross-validation results with the standard deviation to alleviate the bias from the data split.

### 5.1 Task 1: 3D Neuron Detection in 3D NeuroPAL Volumes

#### Quantitative results

We evaluated pre-trained amd retrained detection models, establishing a baseline for comparison in Tab. 2. CellPose used two-channel images to match its pretraining setup, achieving better out-of-the-box performance than pretrained Micro-SAM. nn-UNet outperformed all models, highlighting the effectiveness of fine tuning point-based method for neuron detection.

**Table 2.** Neuron detection results (Task 1). We report metrics using two different distance thresholds for for both pre-trained and re-trained (^†^) models.

| Method | $d_{th}=3\mu m$ | | | $d_{th}=6\mu m$ | | |
| --- | --- | --- | --- | --- | --- | --- |
|  | Precision | Recall | F1 | Precision | Recall | F1 |
| CellPose [18] | $0.45\pm0.02$ | $0.58\pm0.01$ | $0.47\pm0.01$ | $0.50\pm0.03$ | $0.66\pm0.01$ | $0.53\pm0.02$ |
| Micro-SAM [2] | $0.41\pm0.01$ | $0.43\pm0.02$ | $0.39\pm0.01$ | $0.50\pm0.01$ | $0.53\pm0.03$ | $0.47\pm0.01$ |
| nn-UNet<sup>†</sup> [6] | **$0.75\pm0.03$** | **$0.70\pm0.02$** | **$0.71\pm0.02$** | **$0.80\pm0.03$** | **$0.75\pm0.02$** | **$0.76\pm0.01$** |

#### Qualitative results

As shown in Fig. 2 , most errors occur in regions with densely packed neurons of similar colors. The false negative errors in yellow are often due to low intensity or contrast of ground truth neurons blending into background noise. The false positive errors in red are often caused by non-neuronal cells or background noise in the image that are misclassified into neurons.

**Fig. 2.**
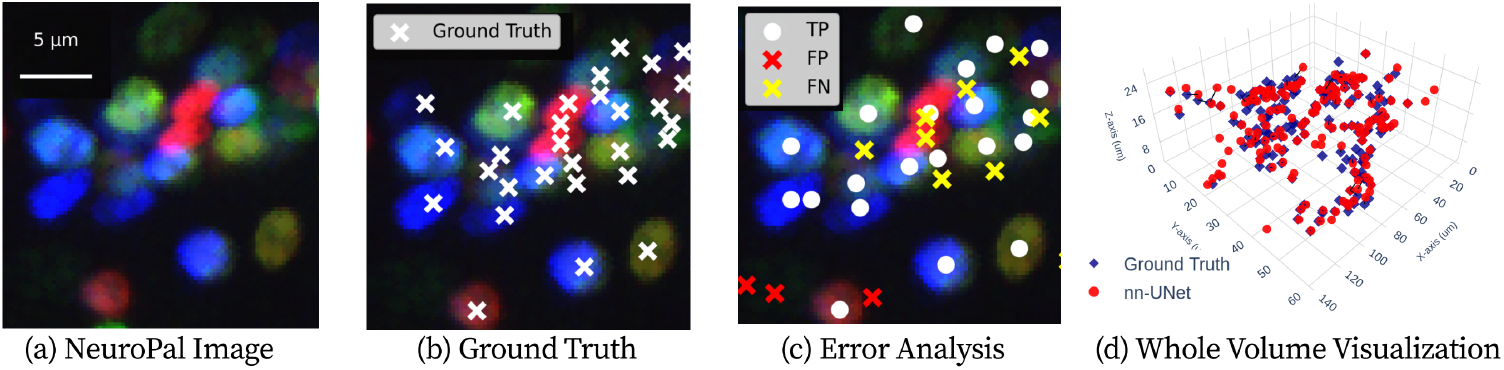
Qualitative neuron detection results (Task 1). (a) A zoomed-in NeuroPAL image, (b) ground truth detection points overlaid, (c) nn-UNet detection results showing true positives, false positives, and false negatives, and (d) whole volume visualization of ground truth and nn-UNet prediction in 3D.

CellPose and Micro-SAM are not natively designed for 3D volumetric NeuroPAL images with color, which presents challenges in directly adapting these methods to such data. Despite this, the pre-trained models produce reasonable results, achieved through the post-hoc stitching of 2D segmentation maps.

### 5.2 Task 2: 3D Neuron Identification in 3D NeuroPAL Volumes

#### Quantitative results

As shown in Tab. 3, all methods have higher accuracy with a low number of labels compared to a high number of labels. Since identification is a classification task, performance improves when there are more candidate choices, which explains the higher accuracy in Top-5 compared to Top-1. The traditional methods (CPD, Statistical Atlas) have relatively lower accuracy than the deep learning methods (CRF ID, fDNC) when they are not finetuned on all datasets. However, there is a significant improvement in their performance after retraining on the full datasets, especially for the CRF ID, which has the best performance among all methods.

**Table 3.**
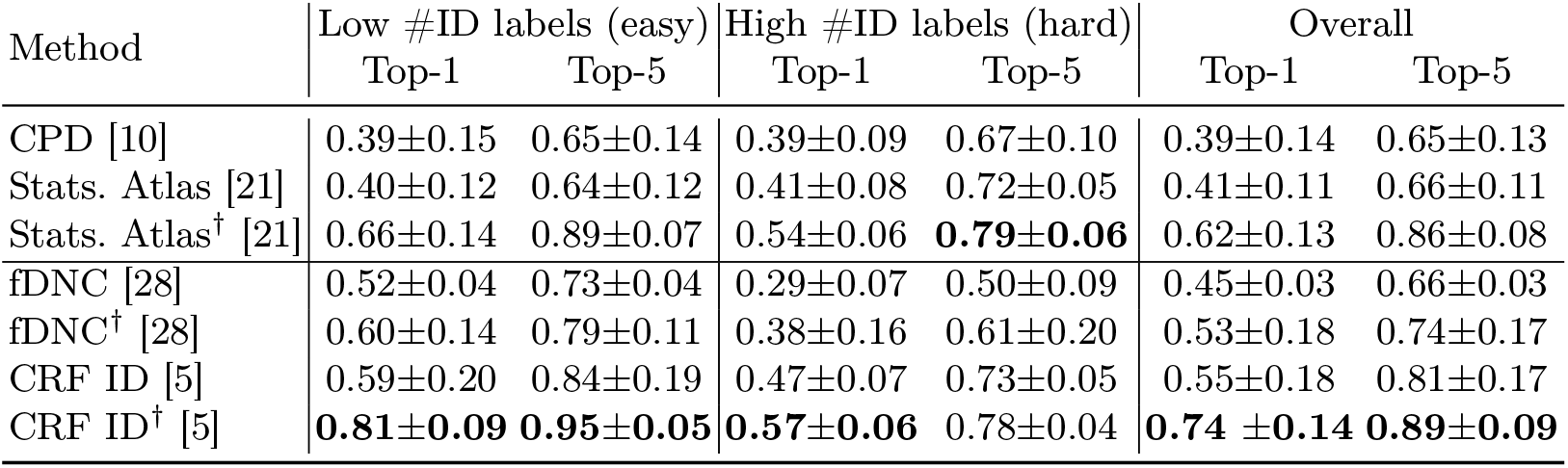
Neuron identification results (Task 2). Mean top-1 and top-5 accuracies are reported for both pre-trained and re-trained (^†^) models.

#### Qualitative results

As shown in Fig. 3, a common source of errors is misclassification due to nearby neurons with similar colors, particularly in methods that rely solely on positional information. The CPD method, which lacks additional information, serves as a baseline for comparison. In contrast, the other three methods improve accuracy by incorporating color and prior probability distributions, enabling more reliable neuron identification. The fDNC method applys a transformer network for matching but is sensitive to worm body orientation during pre-processing, likely contributing to its lower accuracy. The Statistical Atlas method uses a statistical model, while the CRF ID method utilizes a graphical model, leveraging structured prior knowledge for improved performance. The high accuracy of both methods after re-training demonstrates the importance of training on diverse datasets.

**Fig. 3.**
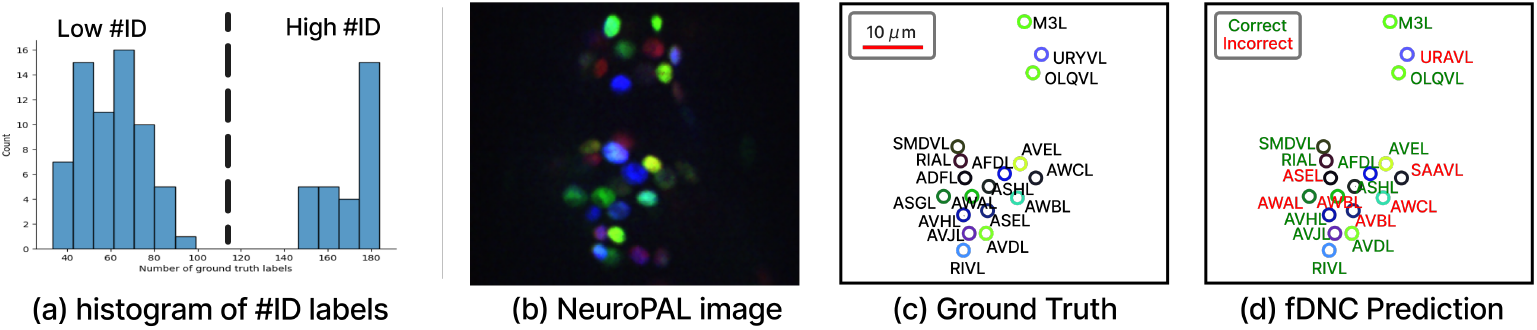
Qualitative neuron identification results (Task 2). (a) We divide the samples into low and high number of ID labels for fine-grained analysis. We show (b) one image from the original 3D NeuroPAL volume, (c) the ground truth label by human experts, and (d) the prediction result by fDNC. The wrong prediction (e.g. ASGL) comes from a nearby neuron with a similar color (AWAL).

### 5.3 Task 3: 3D Neuron Tracking in 3D Calcium Videos

#### Quantitative results

As shown in Table 4, Ultrack consistently outperforms 3DeeCellTracker in both detection (DET) and tracking (TRA) accuracy across distance thresholds. Although both methods use the same StarDist detections, Ultrack benefits from additional ultrametric contours maps. To assess the impact of detection quality on tracking performance, we input ground truth detections at 100%, 80%, and 60% to 3DeeCellTracker. As expected, the DET metric matches the sampling percentage. As shown in Table 5, 3DeeCellTracker significantly improves with better detections, highlighting the need for more accurate detection methods to enhance overall tracking performance.

**Table 4.** Neuron tracking results (Task 3). We report metrics using two different distance thresholds using 5-fold cross-validation.

| Method | $d_{th}=3\mu m$ | | $d_{th}=6\mu m$ | |
| --- | --- | --- | --- | --- |
|  | DET | TRA | DET | TRA |
| 3DeeCellTracker [23] | $0.38\pm 0.06$ | $0.36\pm 0.06$ | $0.42\pm 0.08$ | $0.40\pm 0.07$ |
| Ultrack [4] | **$0.70\pm 0.04$** | **$0.66\pm 0.05$** | **$0.81\pm 0.05$** | **$0.78\pm 0.05$** |

**Table 5.** Tracking results with sampled oracle detections with [23] at *d*_th_=3*μm*.

| Oracle % | TRA |
| --- | --- |
| 100 | 0.99 |
| 80 | 0.74 |
| 60 | 0.52 |

## 6 Limitation and Discussion

WormID-Bench provides a structured framework for benchmarking whole-brain neural dynamics extraction in *C. elegans*, yet several limitations persist. Despite its diverse dataset, it may not fully encompass the range of experimental conditions and imaging setups used across different research groups, potentially introducing domain gaps that affect model generalization. Additionally, existing detection, identification, and tracking methods exhibit inconsistent performance across setups, highlighting the need for more robust, domain-adaptive approaches. Another critical challenge is the cascading error propagation across tasks—errors in neuron detection can lead to misidentifications, which subsequently compromise tracking performance, amplifying inaccuracies at each stage. Future work should explore end-to-end learning pipelines that mitigate these dependencies through joint optimization strategies. Furthermore, while the benchmark promotes standardized evaluation, enhancements such as active learning for dataset expansion, the integration of spatial-temporal priors, and leveraging foundation models could further improve generalization across laboratories.

## Footnotes

1 Videos in SK1 and SK2 have varying z-resolutions and we refer to [17] for details.

## Notes

https://github.com/focolab/WormND

### Competing Interest Statement

The authors have declared no competing interest.

### Summary of Updates

Make a concise version of the paper

